# Multiscale criticality measures as general-purpose gauges of proper brain function

**DOI:** 10.1101/863431

**Authors:** Tomer Fekete, Hermann Hinrichs, Jacobo Diego Sitt, Hans-Jochen Heinze, Oren Shriki

## Abstract

The brain is universally regarded as a system for processing information. If so, any behavioral or cognitive dysfunction should lend itself to depiction in terms of information processing deficiencies. Information is characterized by recursive, hierarchical complexity. The brain accommodates this complexity by a hierarchy of large/slow and small/fast spatiotemporal loops of activity. Thus, successful information processing hinges upon tightly regulating the spatiotemporal makeup of activity, to optimally match the underlying multiscale delay structure of such hierarchical networks. Reduced capacity for information processing will then be expressed as deviance from this requisite multiscale character of spatiotemporal activity. This deviance is captured by a general family of multiscale criticality measures (MsCr). We applied MsCr to MEG and EEG data in four telling degraded information processing scenarios: disorders of consciousness, mild cognitive impairment, schizophrenia and preictal activity. Consistently with our previous modeling work, MsCr measures systematically varied with information processing capacity. MsCr measures might thus be able to serve as general gauges of information processing capacity and, therefore, as normative measures of brain health.

## INTRODUCTION

Over half a century ago, it was first suggested that brains are information processing systems^2^. Consequently, any brain-related dysfunction, should be characterizable as information processing deficiency. This calls for general-purpose measures of such deficiency in brain activity. Seventy years afterwards, such measures are still found wanting. Here, we propose a family of measures and apply them, as a proof of concept, to EEG and MEG datasets in four scenarios: disorders of consciousness (DOC), mild cognitive impairment (MCI), schizophrenia, and in the pre-ictal state.

In face of several processing constraints, such as limited capacity bottlenecks and transmission delays, the brain must represent information hierarchically^4^. Brains succeed in this through division of labor, in which local networks share similar specializations. However, to ensure coherence in the resulting distributed representation, local networks need to be dynamically synchronized, forming components of a higher order network, coarser in spatial and temporal grain (see figure 1). Successful information processing critically depends on this process of orchestration proceeding recursively up to the level of the entire brain^5–7^. Therefore, during peak operation, neuronal activity must generate spatiotemporal activity reflecting this hierarchical organization of processing. Specifically, activity will possess just the right admixture of a multitude of small-scale fast activity (resulting from local 1^st^ order processing) together with much less frequent increasingly slower and more wide-spread events (resulting from higher order orchestration), in effect matching the hierarchical delay structure embedded in neural network organization. Consequently, deviance from this optimal organization will gradually disrupt information processing until the point it altogether vanishes.

**Figure 1:**
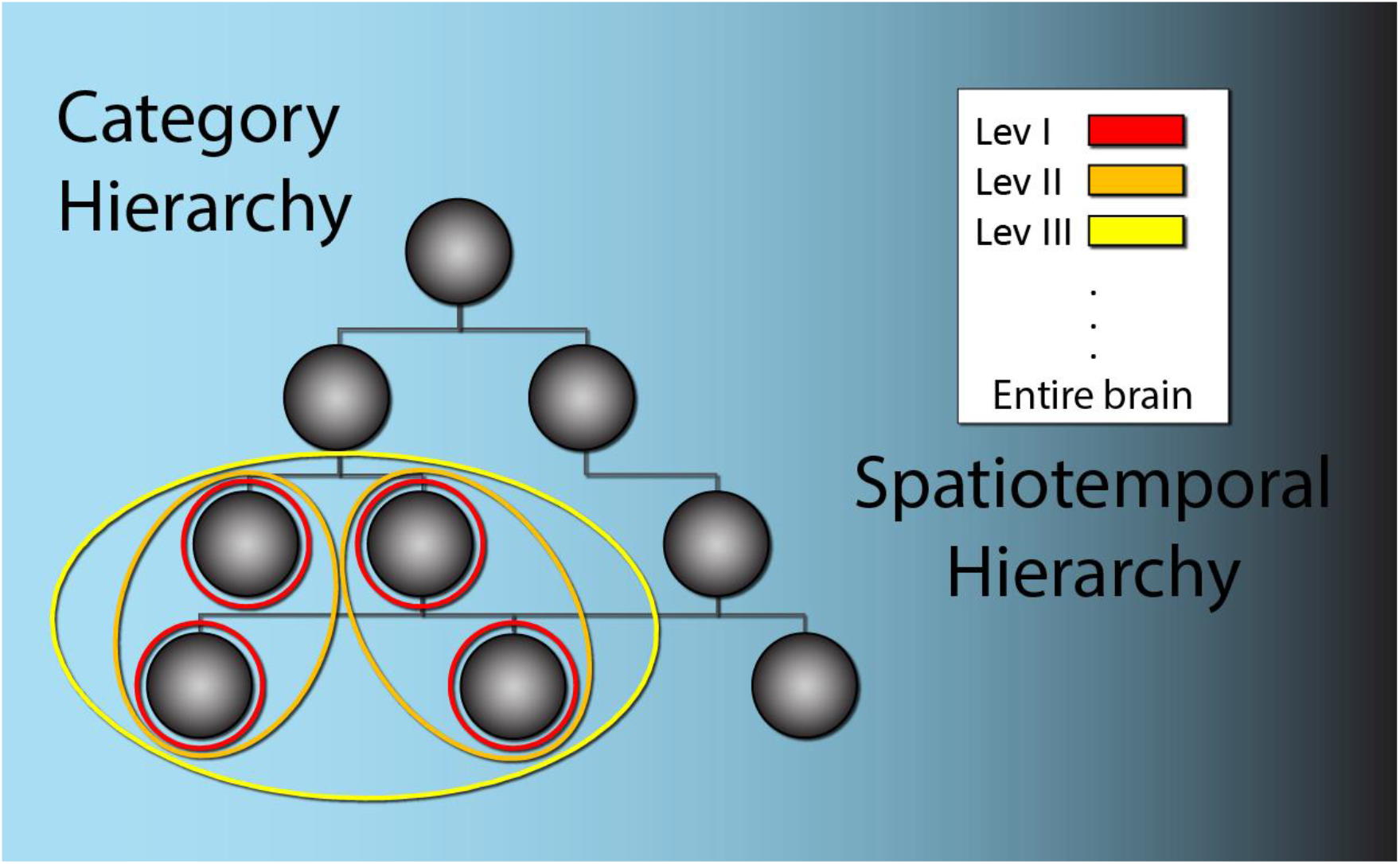
The importance of the hierarchical spatiotemporal structure of neural information processing. Each node in the tree (gray full circles) represents a category, starting from basic features, through objects and ultimately abstract concepts. For illustration purposes we assume each such node is processed by a single local network (red circles). Local networks processing related information need to be orchestrated into a higher order network (“binding”). This process must take place recursively up to the highest level of organization – the entire brain^3^. Because of synaptic delays, which depend on distance, as such higher order networks increase in scale spatially with order, length of communication cycle increases as well. Therefore, activity orchestrated to efficiently match this organization - a necessary precondition for effective information processing– will exhibit the right mixture of fast local events, and increasingly less frequent slower and more extended events. Conversely, the further activity is removed from optimal spatiotemporal makeup, information processing will degrade, until it all but vanishes. The latter can occur naturally, such as during dreamless sleep, as a result of pharmacologically active substances, e.g. anesthetics, or due to brain maladies, e.g. conditions leading to minimally conscious states.

The framework of critical dynamics lends itself to characterization of the spatiotemporal structure of activity. The basic units of analysis are neuronal avalanches^8^ - spatiotemporal bouts of activity - whose basic properties (e.g. size, duration) give rise to distributions. Such distributions tend to be scale free, which is to say, follow a power law^8–21^, a property that is maintained across a wide range of species, conditions and experimental setups (e.g. wakefulness vs. some anesthetic protocols^1^). Thus, criticality analysis results in several power law exponents (see table1), associated with the expression of avalanches in a given state of the brain (in the sense of^22^). Recently^1^ we have shown that, when criticality analysis is applied recursively across several temporal grains, the scale-dependence of these exponents and their pairwise relationships are sensitive to the state of the animal (e.g. awake monkey vs under anesthesia). Using a model of a mesoscopic cortical network, we were able to show that multi-scale criticality (MsCr) measures co-vary systematically with network state/composition (e.g. with excitation-inhibition balance; EIB).

**Table 1:** Notation used to denote MsCr measures. See Fig. 2 for an illustration of computing MsCr features as well as for a detailed description^1^.

|  |  |
| --- | --- |
| $\sigma$: neural gain parameter (branching parameter) | $\gamma$: The exponent of the power law fit to the avalanche average size per duration distribution |
| $\alpha$: The exponent of the power law fit to the avalanche size distribution | $\delta$: The exponent of the power law fit to inter avalanche interval distribution |
| $\tau$: The exponent of the power law fit to the avalanche duration distribution | a, b: the slope and intercept of the linear fit to a given exponent (or pair of such) time dependent curve, e.g. $a_{\sigma,\alpha}$ would be the slope of $\alpha(\Delta t)$ plotted against $\sigma(\Delta t)$ |

Crucially, these measures also co-vary with information theoretic measures (e.g. pattern entropy, pairwise mutual entropy, and Lempel-Ziv complexity, which has been shown to vary with anesthesia level)^23^. MsCr measures could thus serve to determine whether the conditions for neural information processing are optimally realized – namely whether activity is regulated to possess the requisite spatiotemporal makeup. Conversely, MsCr measures will be systematically shifted from optimal values when brain function, and hence information processing, is compromised. To test this hypothesis, we applied MsCr analysis to four EEG and MEG data sets collected from populations in which function is compromised: subjects with disorders of consciousness (DoC), subjects with mild cognitive impairment (MCI), in schizophrenia, and during interictal and preictal activity in epilepsy.

## METHODS

### Data preprocessing

Data were preprocessed separately for each sensor type (EEG, MEG gradiometers and magnetometers) andwere analyzed using a combination of EEGLab routines and custom code. Data were first high-pass filtered (cut-off 1Hz), then a customized adaptive filter was applied to suppress line-noise. This followed by Artifact Subspace Reconstruction^24^, re-referencing to the mean for EEG, and low-pass filtering (cutoff 40Hz). Next Infomax ICA^25^ was carried out. The resulting ICs were evaluated automatically for artifacts, by combining spatial, spectral and temporal analysis of ICs in conjunction with analysis of EOG channels where they were recorded (see below). ICs identified as containing ocular, muscular or cardiac artifacts were removed from the data. For a detailed description of the custom routines, as well as the source code see (Dotan et al, forthcoming).

### Avalanche analysis

Altogether 33 features pertaining to the spatiotemporal scaling behavior of each data set were derived. To extract avalanches from multi-sensor arrays, data need to be coarse-grained (i.e. choosing a time scale), and then discretized using a thresholding operation. Avalanches are defined as clusters of events across channel (i.e. with no interspersing quiet bins across channels). Avalanches were extracted for a range of thresholds (2.5-3.5 stds for EEG data and 3-4 stds for MEG data) and time scales (1-10Δ*t*). Following our previous work, which indicated that avalanche distributions are relatively insensitive to choice of threshold within appropriate ranges^1,19^, distributions were averaged across thresholds to increase signal to noise. The exponents describing the size (α), duration (τ), average size per duration (γ), and inter avalanche interval distributions (δ) were derived using a Maximum Likelihood estimator (the procedure is described in detail in^19^), as well as the branching parameter (σ). Next, each of these 5 parameters was assessed for its temporal scale dependent profile (by deriving the slope and intercept of each time dependent curve – e.g. *a*_*γ*_, *b*_*γ*_ for *γ*(*k*Δ*t*), *k* = 1,…,5). Similarly, the time dependent curves for each pair of these 5 parameters (altogether 10 pairs) were analyzed (deriving slope and intercept; e.g. *a*_*σ,τ*_ and *b*_*σ,τ*_) producing an additional 10×2 parameters. Finally, the magnitude distribution was analyzed for its deviance from an ideal power law using the kappa and generalized kappa measures, and the drop-off of each such distribution was derived.

For each data set, MsCr measures were derived for three intervals, 1-5Δ*t*, 3-7Δ*t* and 6-10Δ*t*. We report the most significant result for each data set, and hence conservatively only p-values smaller than 0.01.

ANOVA/t-test results were corrected for each data set across the 33 MsCr measures using an FDR approach^26^. These results were compared to similar non-parametric tests namely Kruskal–Wallis/Mann–Whitney corrected for multiple comparisons in a similar fashion.

### DOC data

The data comprise 183 EEG records from 4 groups – control healthy subjects and patients emerging from minimal consciousness state, minimally conscious state and vegetative state (VS: n=77, MCS: n=70, EMCS: n=24, control: n=12). EEG was recorded using an EGI 256 electrode sensor net, during a ‘Global-Local’ auditory task^27^. Data were epoched into 1.5 sec segments around stimulus onset (for a complete description of these data and procedure see^28^, with the difference that there only part of each trial was analyzed). EOG channels were defined from the appropriately located electrodes in the sensor net.

### MCI data

Data were collected at the neurology department at the OVGU hospital, Magdeburg, Germany. All procedures were approved by the hospital’s ethical board, and informed consent was obtained from all participants. Data comprise combined EEG/MEG (32 channel/102/204 magnetometers/planar gradiometers Elekta NeuroMag+EOG) records collected from 75 participants (21 controls). There were no significant differences in group demographics: age (average patient age was 68.47±1.06 years, while average control age was 69.07±3.04 years) sex and handedness. During scanning, participants underwent an attention detection task, adapted from a previous task reported in^29^. The task comprised 4 blocks whereby block 1 and 3 presented the attentional task and block 2 and 4 a simple detection task. The stimuli consisted of 4 squares each filled with a single color chosen randomly from 8 possible colors. Stimuli were presented in the center of the screen covering a visual angle of about 2°. Stimuli persisted for 400ms, with ISIs randomly drawn from the 1.3-1.7s range. The total duration of each block was 280s, including a short 8s pause every 40s. In the attentional condition (blocks 1 and 3) participants were asked to indicate the presence of a red colored square among the 4 squares with a button press with their right middle finger of the right hand and otherwise press a button with the right index finger. The probability for the appearance of red in any of the 4 squares was 15%. In the simple detection task (blocks 2 and 4) participants were press a button with the right index finger as soon as the stimulus appeared. Participants were instructed to fixate to the center of the screen and press the buttons as fast and accurately as possible.

For avalanche analyses, the magnetometers channels were concatenated with EEG channels, for a total of 134 channels. Before further analysis, data were down sampled to 250Hz.

### Schizophrenia data

Resting data were acquired during rest using an MEG SQUID sensor array comprising 275 radial first order gradiometers uniformly distributed over the inner surface of a whole-head helmet (600Hz sampling frequency; 0-150 Hz bandwidth; CTF Systems, Inc., Coquitlam BC, Canada).

Data included a total of 100 controls and 56 patients. Control groups included 63 healthy females and 37 males (mean age: 32.47 ± 1.11 years) and patient groups consisted of 16 females, 40 males (mean age: 33.53 ± 1.85 years) diagnosed following DSM-IV criteria. Out of controls, 8 were left-handed. Among the patients, 2 were left-handed. Controls did not have any neurological or psychiatric illnesses or any history of head trauma. All the controls underwent Structured Clinical Interview and were reported normal. Patients were all under anti-psychotic medication.

Before further analysis, data were down sampled to 250Hz. Additionally, data from 2 broken sensors were removed for all subjects.

### Epilepsy data

The data were obtained from the European epilepsy database – EPILEPSIAE^30^. This database contains annotated long-term EEG recordings, obtained with scalp electrodes, from 31 patients. The data consist of 19 channels sampled at 256 Hz. The data were separated into 6235 segments of 4 minutes, with approximately half of the segments marked as pre-ictal and extracted from up to an hour before a seizure, and the other half marked as inter-ictal and extracted from a period a least four hours before/after the occurrence of a seizure.

## RESULTS

We applied multi-scale criticality (MsCr) analyses to 4 different datasets (see Methods) and examined their utility in discriminating between different information processing states of the brain. Below we present the results obtained from each of these datasets in detail. In MsCr, the spatiotemporal behavior of a system (network) is characterized by observing the change in the properties of the spatiotemporal activity patterns it produces – taken as neuronal avalanches – as a function of temporal scale. Figure 1A illustrates the process of deriving MsCr measures from criticality measures iterated across increasing temporal scales. The behavior of such measures as a function of temporal scale is well described by a linear fit. Therefore, the temporal multiscale behavior of given criticality measures can be expressed by two terms – the slope and intercept of the fit.

### MsCr in disorders of consciousness

We reanalyzed a previously published dataset^28^, comprising high density EEG collected during an auditory task from 4 groups: control subjects, emerging from minimal consciousness, minimally conscious and vegetative state. Significant difference among the groups was found for 13 out of the 33 MsCr features (ANOVA, at least *p* < 0.01 corrected for multiple comparisons– see Methods; see table 2, figure 2A as well Methods for feature definition). The basic statistics for each group are given in supp. Table 1. Similar results were obtained also using non-parametric tests (supp. Table 5/6).

**Table 2:**
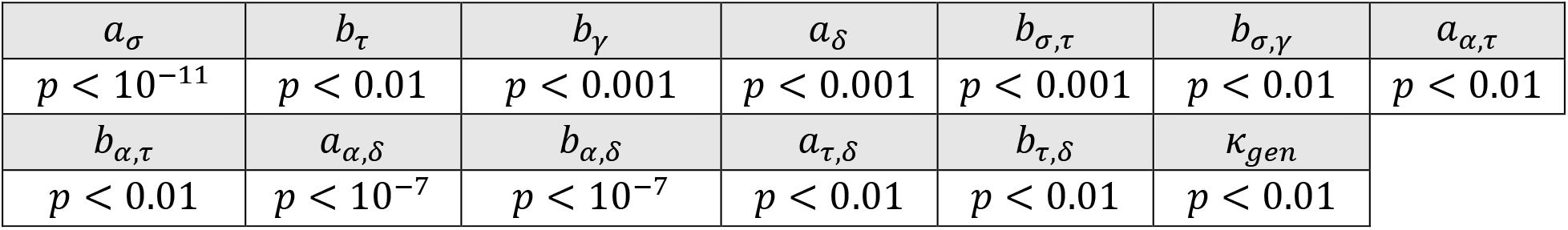
significant MsCr measures in DOC.

|  |  |  |  |  |  |  |
| --- | --- | --- | --- | --- | --- | --- |
| $a_\sigma$ | $b_\tau$ | $b_\gamma$ | $a_\delta$ | $b_{\sigma,\tau}$ | $b_{\sigma,\gamma}$ | $a_{\alpha,\tau}$ |
| $p < 10^{-11}$ | $p < 0.01$ | $p < 0.001$ | $p < 0.001$ | $p < 0.001$ | $p < 0.01$ | $p < 0.01$ |
| $b_{\alpha,\tau}$ | $a_{\alpha,\delta}$ | $b_{\alpha,\delta}$ | $a_{\tau,\delta}$ | $b_{\tau,\delta}$ | $\kappa_{gen}$ | |
| $p < 0.01$ | $p < 10^{-7}$ | $p < 10^{-7}$ | $p < 0.01$ | $p < 0.01$ | $p < 0.01$ | |

**Figure 2:**
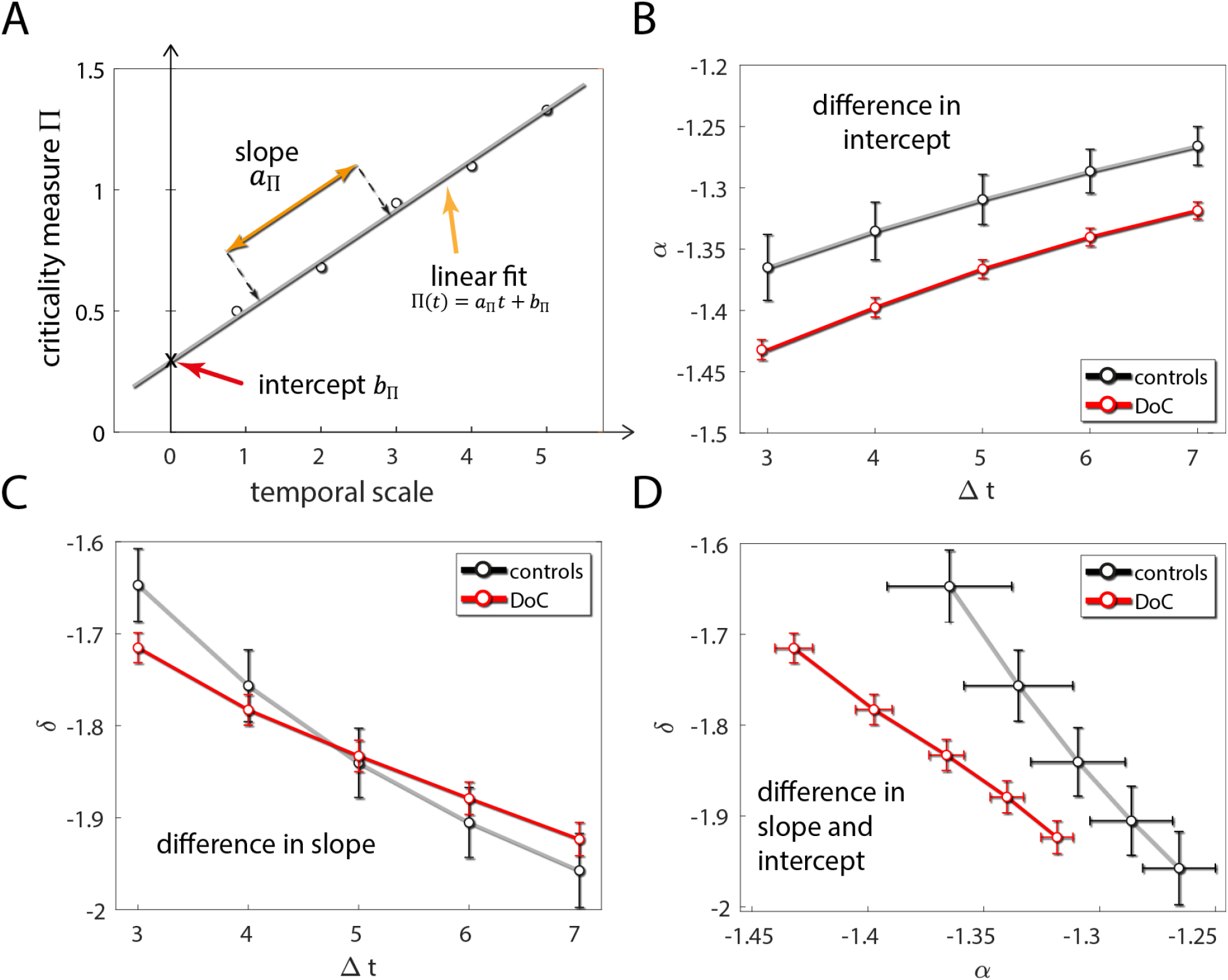
deriving MsCr measures from standard criticality measures. (A) an illustration of deriving MsCr measures. Exponents characterizing a given aspect of the distribution of neuronal avalanches in an experimental condition (e.g. of the distribution of avalanche sizes during sleep) are computed across several temporal scales. They are then fit with a line, and the slope and intercept are taken to represent the multiscale temporal behavior of a subject in a given state of the brain.(B) Difference in experimental condition can manifest in difference in intercept – in this case between subjects suffering from disorders of consciousness (see details below) and controls for the avalanche size distribution (C) Difference in experimental condition can manifest in difference in slope - here for the inter-avalanche interval (IAI) across temporal scale (same data as B) (D) Difference can also manifest in both slope and intercept – in this case in the joint behavior (interaction) of avalanche size and IAI when temporal scale increases.

### MsCr in mild cognitive impairment

MsCr measures were derived from joint MEG/EEG collected during an attention task (see METHODS). Significant difference between MCI patients and matched controls (two-sided t-test; at least *p* < 0.01 corrected – see Methods) was found for 20 out of the 33 MsCr features (see table 3, figure 3B). The basic statistics for each group are given in supp. Table 2. Similar results were obtained also using non-parametric tests (supp. Table 5/6).

**Table 3:** significant MsCr measures in mild cognitive impairment (MCI).

|  |  |  |  |  |
| --- | --- | --- | --- | --- |
| $b_\sigma$ | $a_\alpha$ | $b_\alpha$ | $a_\tau$ | $a_\gamma$ |
| $p < 0.001$ | $p < 0.001$ | $p < 0.001$ | $p < 10^{-4}$ | $p < 0.001$ |
| $b_\delta$ | $a_{\sigma,\alpha}$ | $b_{\sigma,\alpha}$ | $a_{\sigma,\tau}$ | $b_{\sigma,\tau}$ |
| $p < 0.01$ | $p < 10^{-5}$ | $p < 10^{-6}$ | $p < 10^{-4}$ | $p < 0.001$ |
| $a_{\sigma,\gamma}$ | $b_{\sigma,\gamma}$ | $a_{\sigma,\delta}$ | $b_{\sigma,\delta}$ | $a_{\alpha,\tau}$ |
| $p < 10^{-4}$ | $p < 0.001$ | $p < 10^{-4}$ | $p < 0.001$ | $p < 10^{-4}$ |
| $b_{\alpha,\tau}$ | $a_{\tau,\delta}$ | $b_{\tau,\delta}$ | $b_{\gamma,\delta}$ | $\kappa_{gen}$ |
| $p < 10^{-4}$ | $p < 10^{-4}$ | $p < 10^{-5}$ | $p < 0.01$ | $p < 0.001$ |

**Figure 3:**
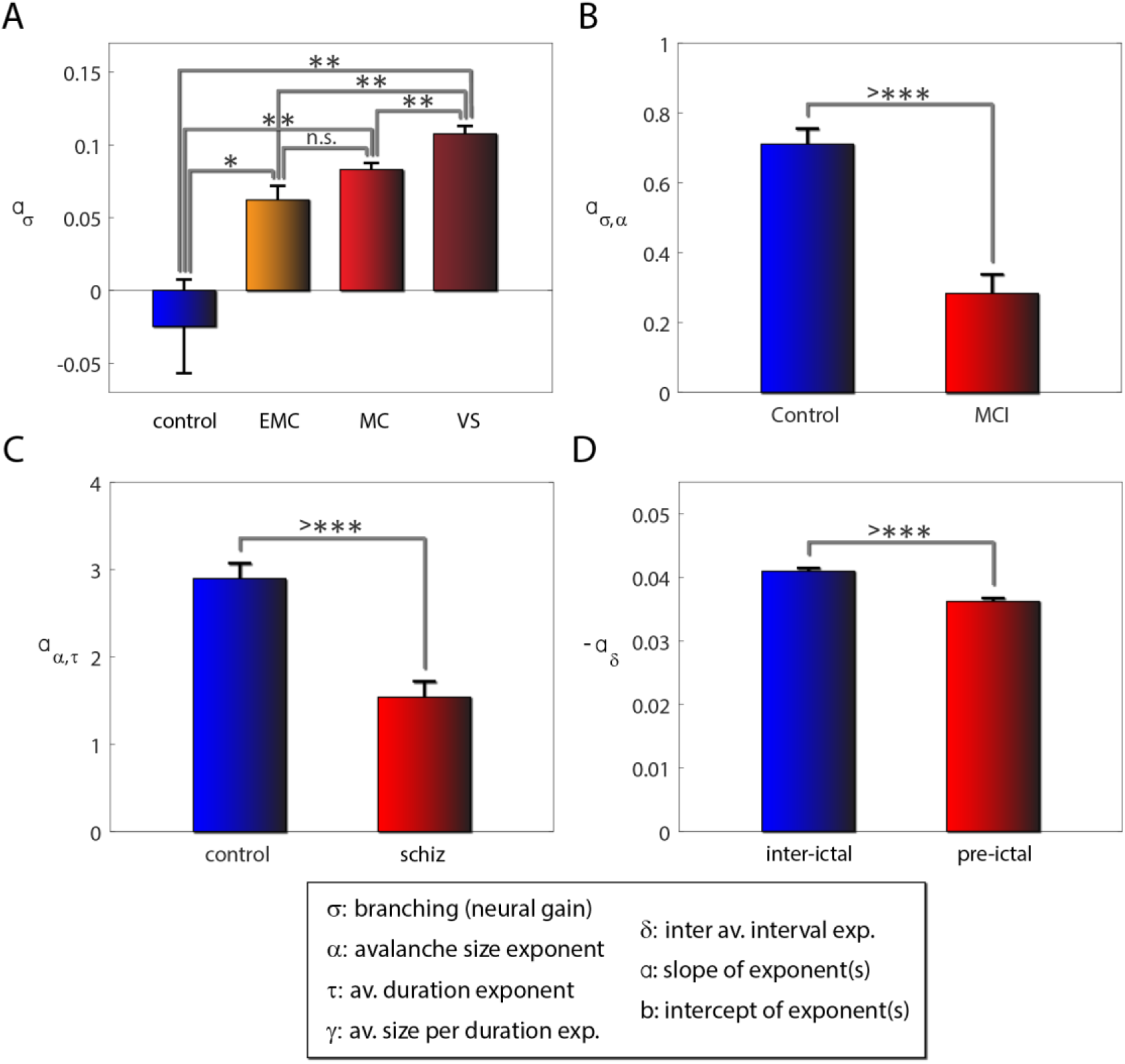
MsCr measures are sensitive to compromised capacity for information processing. The most significant MsCr features for each of the analyzed data sets. (A) The time dependent behavior of the neural gain parameter σ differentiates between control subjects and DOC patients and between vegetative state (VS) and minimally conscious (MC) subjects. MsCr measures were derived from high density EEG collected during an auditory task (B) The slope of the σ α curve a_σ,α_, derived from joint MEG/EEG collected during an attention task is attenuated in MCI patients. This curve describes the time dependent relationship between the neural gain parameter σ, and the exponent α of the avalanche size distribution. (C) The slope of the α δ curve a_α,δ_, derived from resting state MEG is attenuated participants with schizophrenia. This curve describes the time dependent relationship between the exponent α of the avalanche size distribution, and the exponent δ, of the inter-avalanche interval distribution (IAI). (D) The slope of the δ curve a_δ_, derived from resting state EEG is attenuated during pre-ictal activity. This curve describes the time dependent behavior of the IAI distribution.

**Figure 4:**
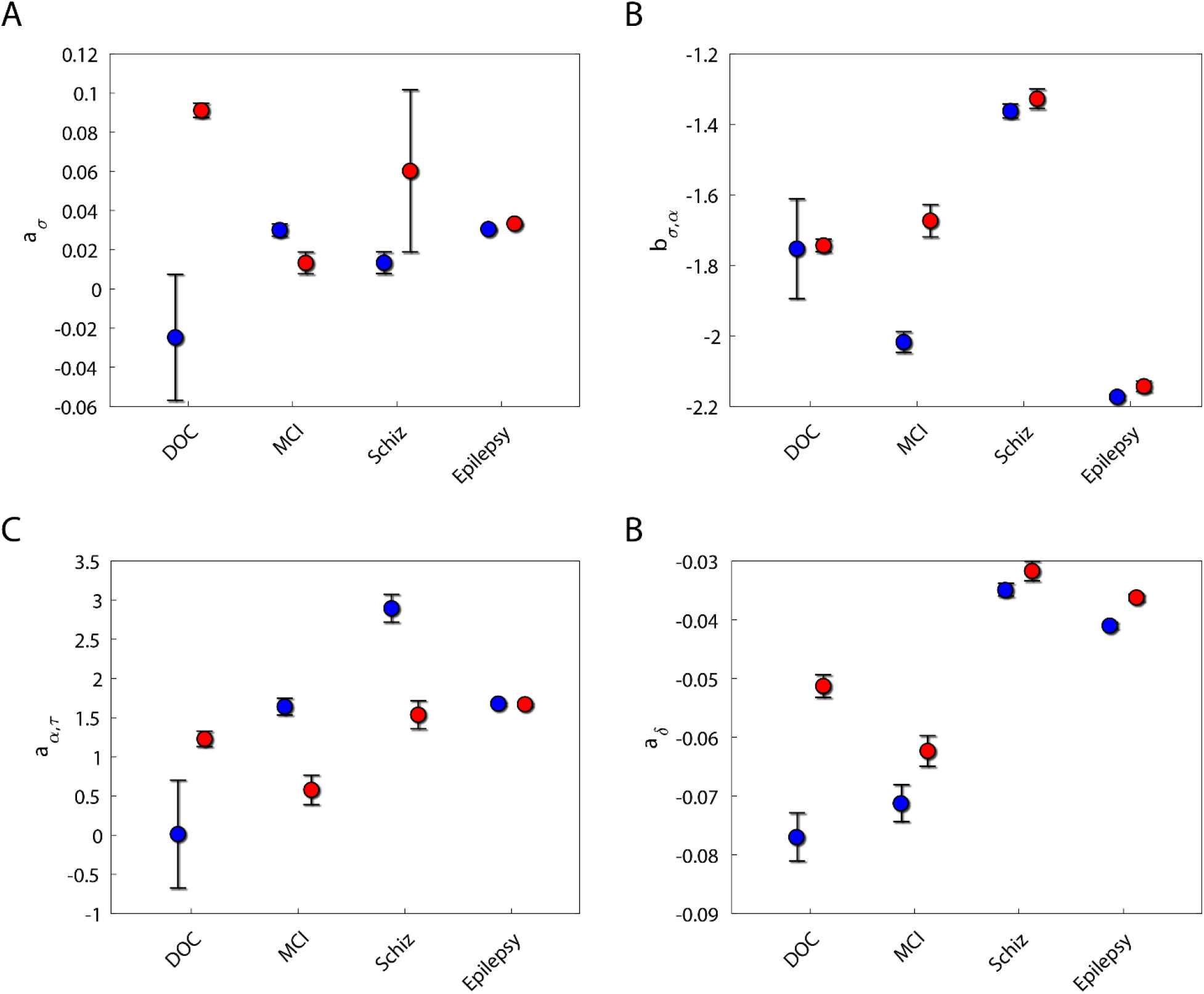
The most significant features in each dataset (see Fig. 3) across all datasets. In general, MsCr features displayed disparate patterns across the different populations. This is consistent with previous modeling work showing that MsCr measures are tuned differentially to different network information processing characteristics. This implies that it is the co-variation of MsCr measures, rather than the values of a single such measure, that in fact characterize information processing states of the brain. Note that MsCr measures, as multiscale measures are sensitive not only to temporal grain but also to spatial grain. The datasets we analyzed were collected from arrays differing in their temporal resolution and sensor density, therefore accounting for (some of the differences) in control population values.

### MsCr in Schizophrenia

MsCr measures were derived from resting state MEG collected from schizophrenia patients and healthy participants. Significant difference between patients and controls (two-sided t-test; at least *p* < 0.01 corrected – see Methods) was found for 8 out of the 33 MsCr features (see table 4, figure 3C). The basic statistics for each group are given in supp. Table 3. Similar results were obtained also using non-parametric tests (supp. Table 5/6).

**Table 4:** significant MsCr measures in Schizophrenia – full data set.

| $a_\tau$ | $a_\gamma$ | $b_\delta$ | $a_{\alpha,\tau}$ |
| --- | --- | --- | --- |
| $p < 0.001$ | $p < 0.001$ | $p < 10^{-5}$ | $p < 10^{-5}$ |
| $a_{\alpha,\gamma}$ | $b_{\alpha,\gamma}$ | $b_{\alpha,\tau}$ | $b_{\alpha,\delta}$ |
| $p < 0.001$ | $p < 0.001$ | $p < 10^{-5}$ | $p < 0.01$ |

Sex was unbalanced between groups. Two-way ANOVA, however, did not find any significant effects of sex, or interactions with MsCr effects. Moreover, discarding controls until balance was obtained resulted in a similar pattern of results.

**Table 5:**
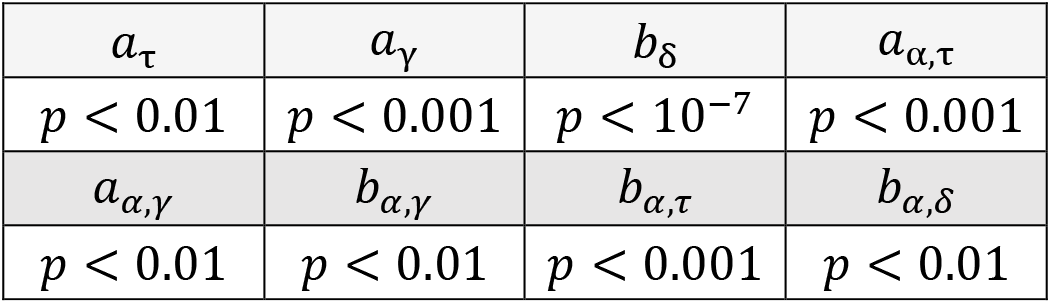
significant MsCr measures in Schizophrenia – balanced dataset. Discarding patients to balance sex between groups doesn’t affect overall pattern of results (compare to balance sex between groups doesn’t affect overall pattern of results (compare to table 4).

|  |  |  |  |
| --- | --- | --- | --- |
| $a_\tau$ | $a_\gamma$ | $b_\delta$ | $a_{\alpha,\tau}$ |
| $p < 0.01$ | $p < 0.001$ | $p < 10^{-7}$ | $p < 0.001$ |
| $a_{\alpha,\gamma}$ | $b_{\alpha,\gamma}$ | $b_{\alpha,\tau}$ | $b_{\alpha,\delta}$ |
| $p < 0.01$ | $p < 0.01$ | $p < 0.001$ | $p < 0.01$ |

### MsCr in Epilepsy

MsCr measures were derived from resting state EEG. Significant difference (two-sided t-test at least *p* < 0.01 corrected – see Methods) between resting EEG recorded during inter ictal activity (at least 4 hours from a seizure) and pre-ictal activity (recorded during the hour preceding a seizure) was found for 14 out of the 33 MsCr features (see table 6, figure 3D). The basic statistics for each group are given in supp. Table 4. Similar results were obtained also using non-parametric tests (supp. Table 5/6).

**Table 6:** significant MsCr measures during pre-ictal activity.

|  |  |  |  |  |  |  |
| --- | --- | --- | --- | --- | --- | --- |
| $a_\sigma$ | $b_\sigma$ | $b_\tau$ | $b_\gamma$ | $a_\delta$ | $a_{\sigma,\tau}$ | $b_{\sigma,\tau}$ |
| $p < 10^{-8}$ | $p < 0.001$ | $p < 0.001$ | $p < 10^{-6}$ | $p < 10^{-8}$ | $p < 0.001$ | $p < 0.001$ |
| $a_{\sigma,\gamma}$ | $b_{\sigma,\gamma}$ | $a_{\sigma,\delta}$ | $a_{\alpha,\delta}$ | $b_{\alpha,\delta}$ | $\kappa$ | $\alpha cutoff$ |
| $p < 0.001$ | $p < 0.001$ | $p < 0.01$ | $p < 10^{-5}$ | $p < 10^{-5}$ | $p < 10^{-4}$ | $p < 0.001$ |

While some MsCr features were highly indicative for some of the datasets analyzed here it was not the case for all of them. Figure 5 portrays the same measures represented in figure 3 across all experiments. Only for one of these features, *a*_*δ*_, the same trend (although not significance) was observed across all four data sets. It is true, that some of the differences between data sets stem from the nature of MsCr measures – as multiscale measures they are also sensitive to spatial scale and not only temporal scale. And indeed, our data derived from different recording apparatus differing in their spatial resolution and sensor density. Nevertheless, for the most part the 33 MsCr behaved differently across datasets (and see also supp. Tables 1–6). This is to be expected, as the cognitive deficits associated with each condition are not one and the same, and as our previous modeling work shows, different facets of information processing co-vary with different patterns of MsCr measures^1^, suggesting that MsCr should be best considered as dynamical fingerprints characterizing different states of the brain.

**Figure 5:**
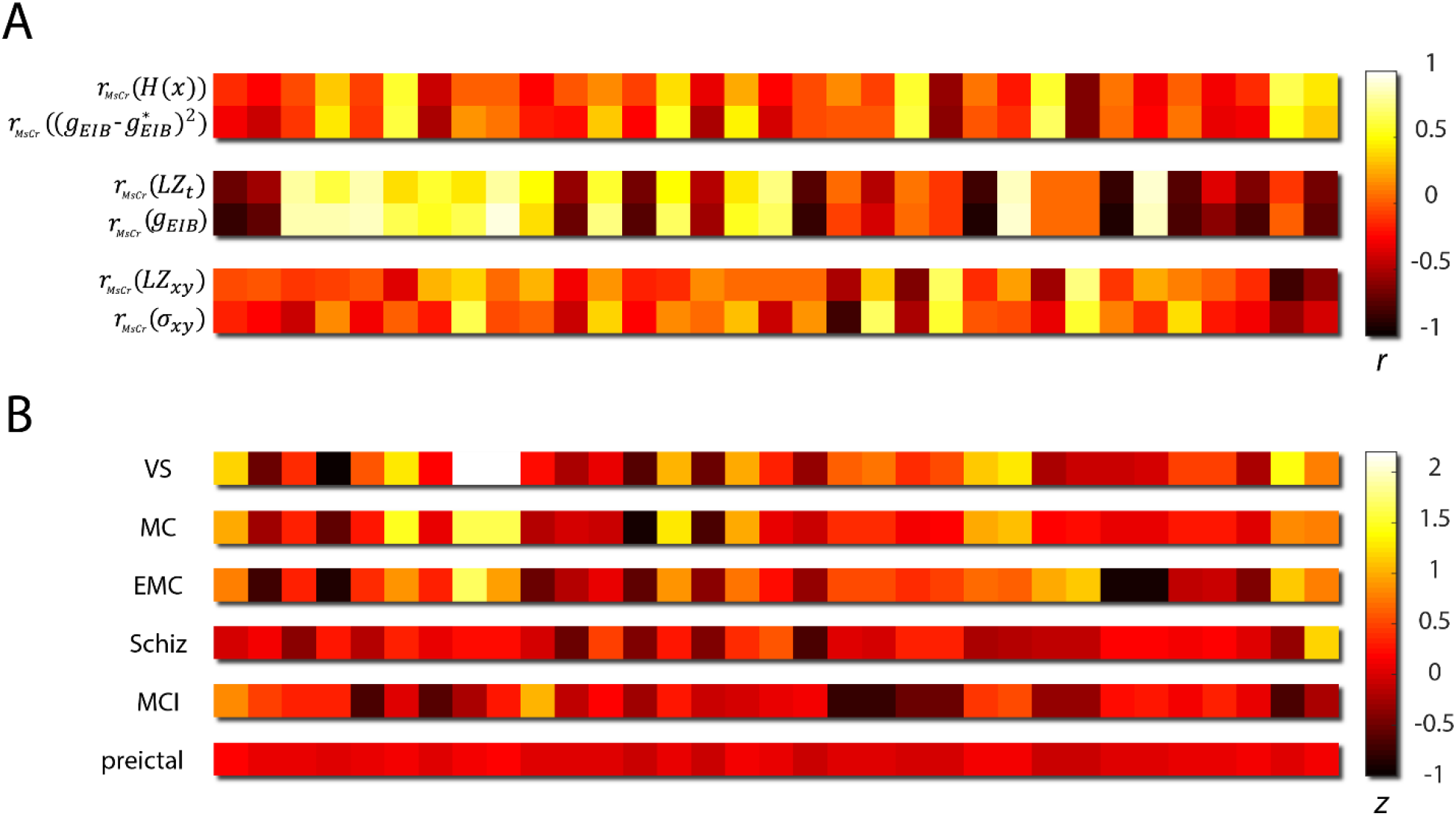
MsCr fingerprints characterize information processing states of the brain. (A) the correspondence between patterns of MsCr correlation to network control parameters, and patterns of correlation between MsCr measures and information measures, in a simulation of a generic cortical network described in^1^. Each horizontal color bar corresponds to a correlation pattern between the 33 MsCr measures used in this study and a variable of interest. The patterns are color coded according to the strength of correlation. r_MsCr_(x) denotes the correlation of x to MsCr measures across various realizations of the simulated architecture. g_EIB_ is the parameter controlling EIB (its increase increases excitation relative to inhibition), 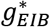 is a numerically derived intermediate value in which informational measures are maximized. σ_xy_ is the parameter used in the simulation to control the spatial extent of connectivity. H(x) denotes activity pattern entropy. LZt and LZxy denote Lempel-ziv temporal and spatial complexity respectively. The correlation between pairs of patterns from top to bottom is 0.95, 0.99 and 0.75 respectively. (B) MsCr fingerprints of compromised informational states of the brain. Magnitude is expressed in z scores relative to the MsCr fingerprint of matched controls. Note that none of the 33 measures exhibits the same relation to the norm (e.g. exceedingly high) across all 6 experimental conditions (see also Fig.4, and supplementary tables 1–4).

## DISCUSSION

### MsCr fingerprints of informational processing states

We set out to investigate whether information processing impairment manifests itself as deviance from typical neuronal activity. Typical activity was depicted in terms of the multiscale spatiotemporal makeup (dynamical fingerprint) associated with alert wakefulness in healthy brains. Upholding this state was considered a necessary condition for achieving meaningful dissemination of information throughout brain networks at various spatiotemporal scales, and hence for information processing at large. Therefore, deviance from this state will characterize the dynamic fingerprints of cognitive and behavioral deficits. To test this hypothesis, we analyzed electrophysiological data, either MEG or EEG, from patients showing a range of deficits.

To capture its dynamical fingerprints, spatiotemporal brain activity was depicted as neuronal avalanches. Statistical properties of their fundamental quantities (e.g. their size and duration distributions) were computed across a range of temporal scales, resulting in multiscale criticality (MsCr) measures. Our previous modeling work^1^ had suggested that MsCr measures are strongly associated with the information processing capacity of generic cortical-like networks, while also varying systematically with the state of the network (e.g. EIB) and its composition (connectivity pattern) – see figure 5a.

As we expected, MsCr fingerprints showed deviance in states of compromised information processing such as disorders of consciousness, mild cognitive impairment, and schizophrenia and even during pre-ictal activity (where mild cognitive deficits such as confusion are common, but not universal^31–33^). Figure 5b represents the MsCr fingerprints associated with the informationally compromised states of the brain examined in this study, in terms of deviance from the MsCr fingerprints of matched controls.

### Relationship to conventional criticality theory

Observations of 1/f scaling in neuronal avalanches have traditionally been understood in terms of criticality, namely the idea that cortical dynamics is poised at the transition between qualitatively different types of dynamical behavior ^34^. Operating near the critical state offers several computational advantages ^18,35,36^, such as increased sensitivity to input, facilitation of information transfer, and increased pattern repertoire. However, there are good reasons to think that the brain is not operating as a critical system, at least not at the macroscopic level. First, criticality involves fine-tuning of the system state, which is antithetic to robustness (say to injury). Moreover, as there is some evidence to the effect that the abovementioned computational benefits are maintained (and perhaps even maximized) in slightly subcritical states of the brain ^6,1,37^. It is therefore unclear what would motivate expending the resources necessary to maintain a critical state. Indeed, while brain dynamics share many features of critical dynamics, nevertheless at no point do they manifest all the features that criticality theory predicts (e.g.^9^ and in contrast to the predictions in ^1^). This led us to suggest that brain dynamics are regulated by various homeostatic processes to be near-critical, that is, give rise to neuronal avalanches that share some (and at times many), though not all, of the scaling properties predicted by theory. This can be phrased in terms of states of the brain (dynamical regimes) lying on a near-critical manifold - a landscape consisting of points each of which represents a state of the brain, in which neuronal avalanches exhibit near critical behavior to varying extents. Alert wakefulness comprises an optimal zone (as far as information processing capacity goes; figure 6a) on this manifold, and to the extent a system moves (e.g. drowsiness) or is pushed away from this zone (e.g. via pharmacology or injury), information processing is increasingly compromised, until it all but vanishes.

**Figure 6:**
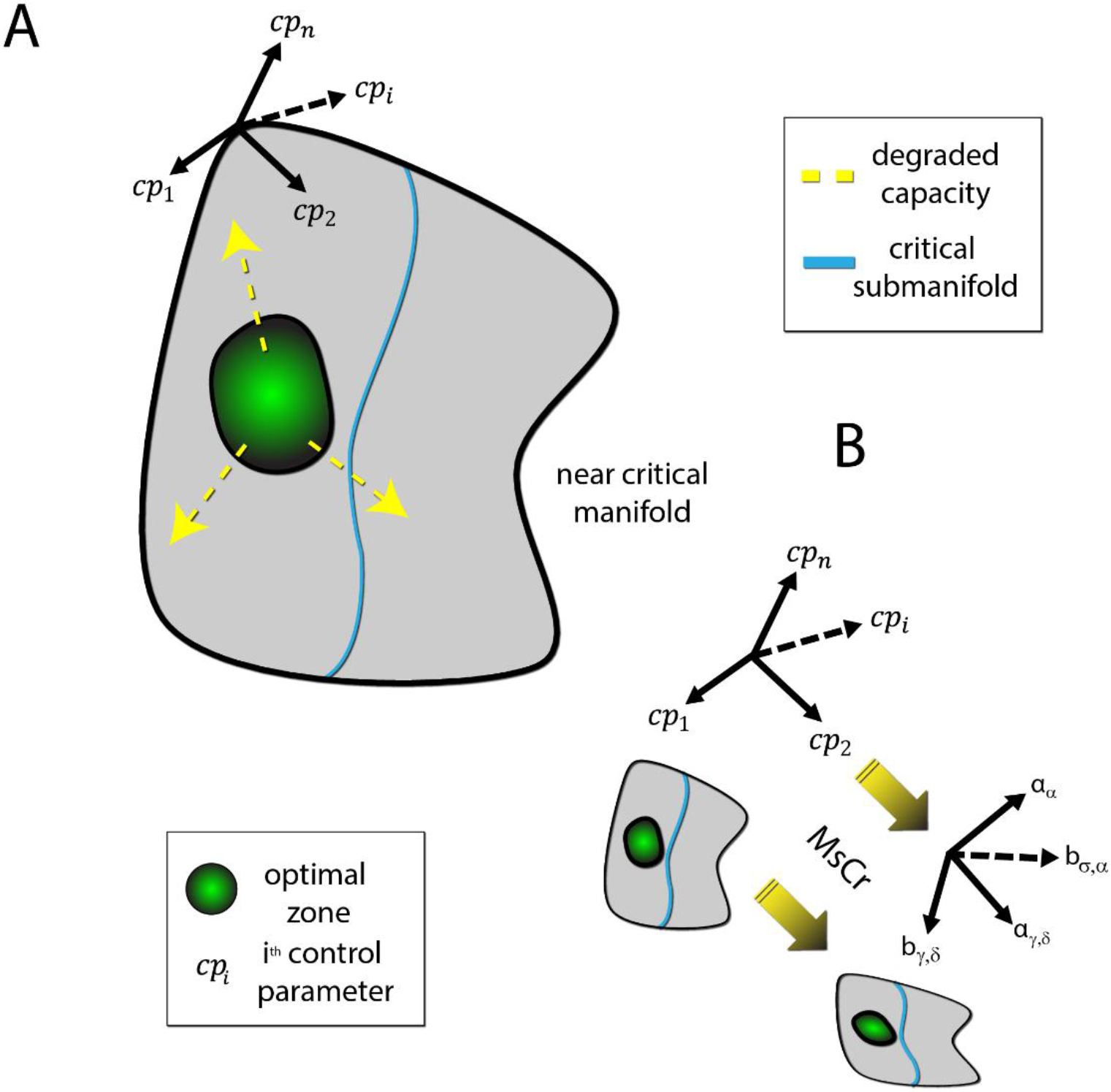
MsCr parameters as bridging constructs between network composition and macroscopic behavior. (A) Healthy alert brains are optimally situated with respect to information processing in terms of the “control parameters” governing neural network function (e.g. EIB, various neuromodulator concentrations, connectivity features). Shift in the information processing state of the brain (from minor changes brought about by drowsiness to profound changes under anesthesia or maladies such as schizophrenia) is brought about by change to network function as a result of change to these parameters (e.g. through neuromodulator concentrations, pharmacology, trauma). (B) Deriving MsCr measures embeds a system’s activity into a new space parametrized by these measures. As our modeling work and previous results suggest, this embedding preserves important information about the topology of the original system’s phase space, which is parametrized by the above-mentioned control parameters. This can be utilized to further the understanding of brain maladies: first the macroscopic deviance in neural activity characteristic of a given condition can be characterized through MsCr analysis – the MsCr “fingerprint”. Next, models such as those suggested in^1^ can be elaborated with detailed network composition features to determine which network properties are the likely culprits. In turn, experiments can select between alternative targets that can reproduce the MsCr fingerprint and help refine modeling. This can also potentially enable rational development and screening of novel therapeutics.

This near-critical manifold of states of the brain is best thought of as parametrized by various “control parameters” – factors which have widespread effects on the dynamics (and hence function) of brain networks. Among those factors would be network properties such as EIB, levels of neuromodulators, and network connectivity. The brain’s ability to regulate such properties allows it, on the one hand, to effectively shut itself off (e.g. during sleep) but at any point with relative ease switch back sufficiently close to the optimal zone when sudden alertness is needed, for instance to engage in fight or flight behavior.

From this perspective, deriving MsCr fingerprints can be viewed as a transformation, embedding the near critical manifold in a new space parametrized by MsCr measures (figure 6b). Our results indicate that this transformation preserves important features of the topology of this manifold. This suggests that MsCr can serve as a mechanism linking between collective behavior (at the macro or meso scales) and underlying network structure. To the extent this idea holds, MsCr (or in the future more exact multiscale complexity measures deriving from neural computational theories), can cast the illusive ideas of ‘function’ and ‘information processing’ in operational terms, affording significant clinical benefits. For example, better understanding of states of the brain in which information processing is compromised (and neuronal systemic health) would follow from an iterative “procedure”: first, MsCr fingerprints of such states of the brain can be identified. Next, modeling can capitalize on these empirical generalizations by exploring which network properties can give rise to them. Experimentation at the sub-network level in animal models can then select between competing potential mechanisms, and in turn help refine modeling. This invokes an intriguing prospect – namely employing multiscale complexity measures such as MsCr to try and coax maligned network dynamics (say due to a neurological disorder, or after trauma) towards health. This would be achieved through employing MsCr fingerprints of healthy brain as targets to be attained via pharmacology and brain stimulation, potentially facilitating rational development of therapeutics.

## ACKNOWLEDGEMENTS

We wish to express our gratitude to Cees van Leeuwen and Matteo Colombo for their helpful comments on an earlier version of this manuscript.

## SUPPLEMENTARY INFORMATION

**Supplementary Table 1:**
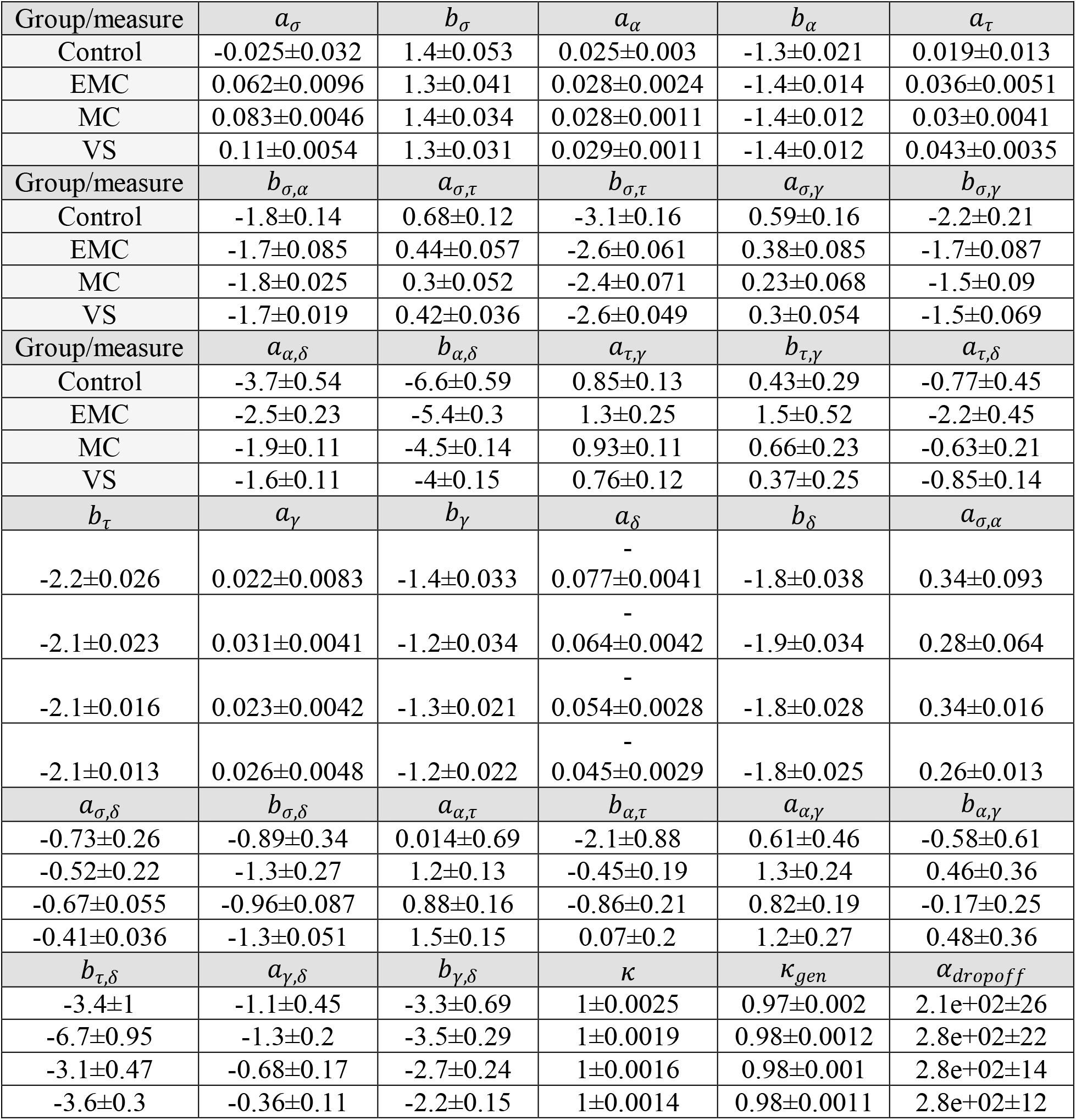
Group averages and standard errors of the 33 MsCr measures used in this study derived from the DOC dataset

**Supplementary Table 2:**
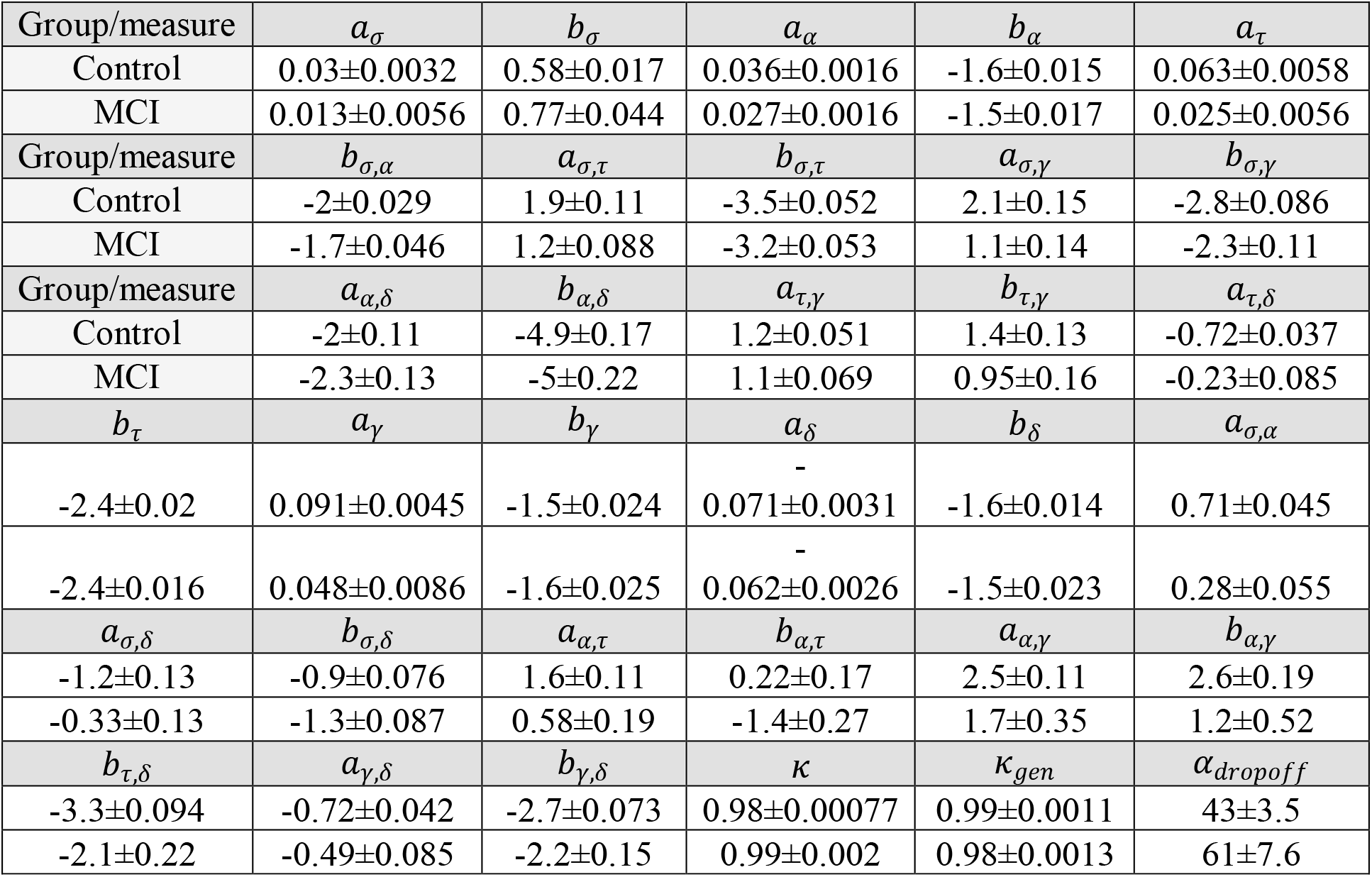
Group averages and standard errors of the 33 MsCr measures used in this study derived from the MCI dataset

**Supplementary Table 3:**
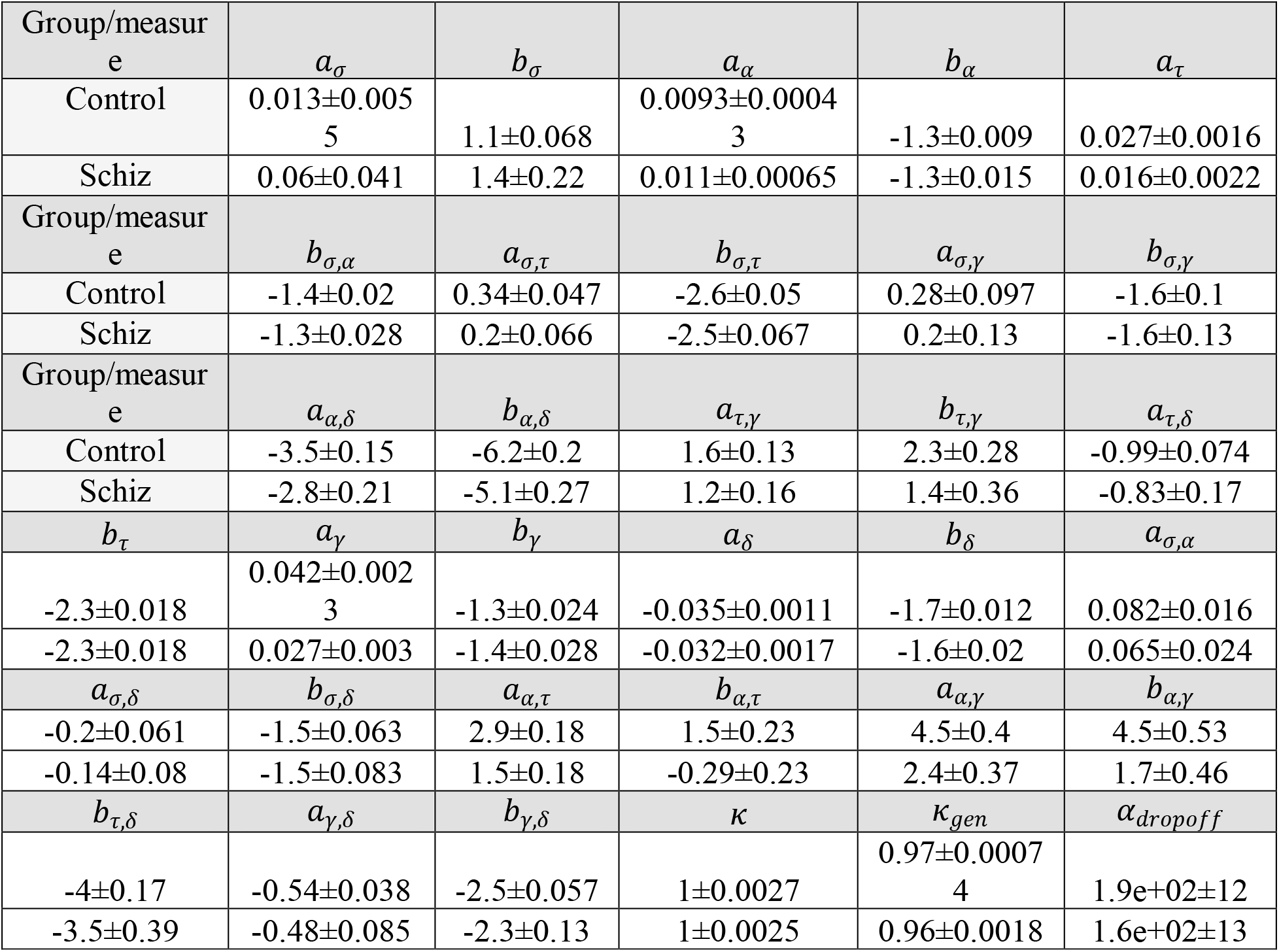
Group averages and standard errors of the 33 MsCr measures used in this study derived from the Schizophrenia dataset.

**Supplementary Table 4:**
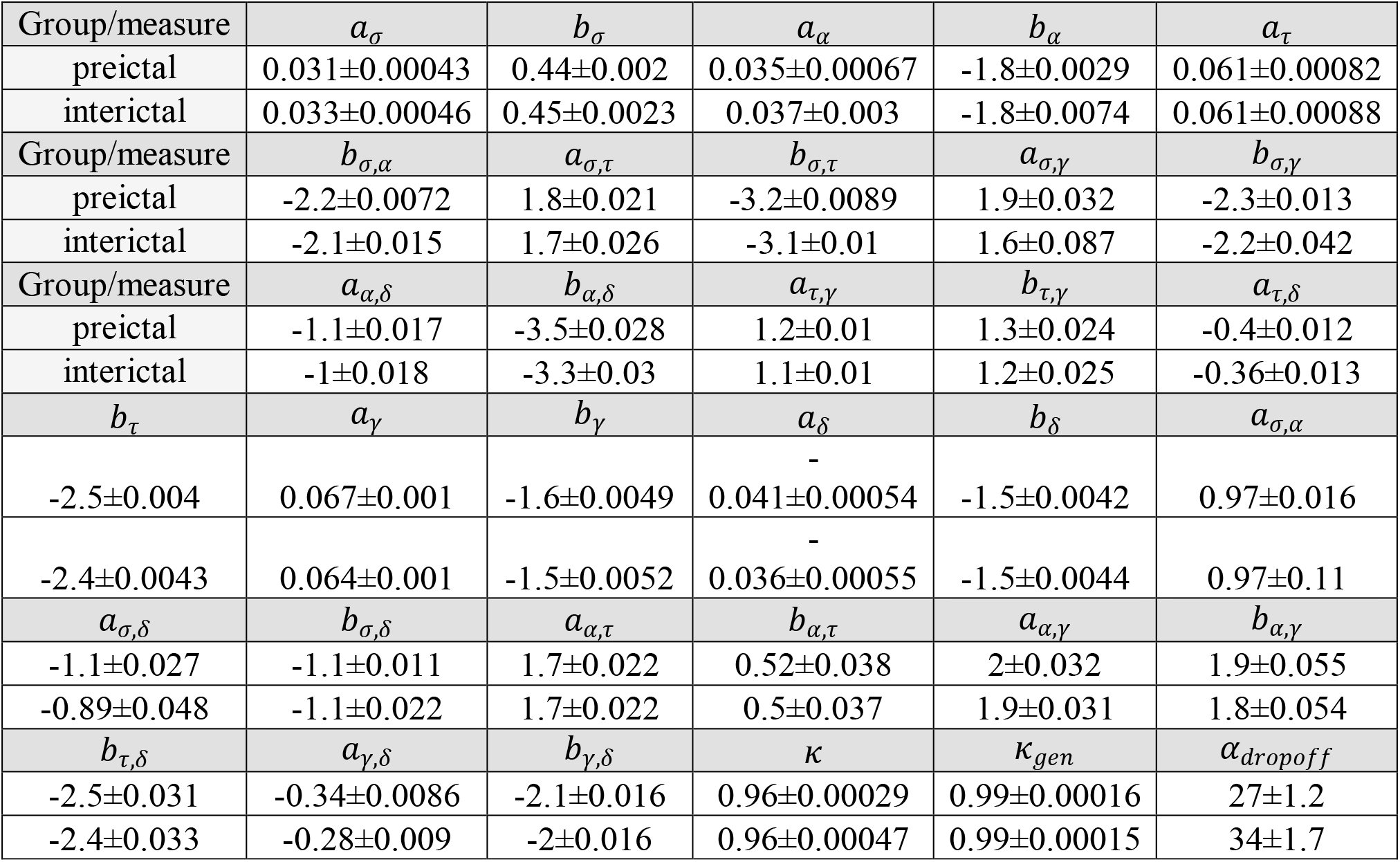
Group averages and standard errors of the 33 MsCr measures used in this study derived from the epilepsy dataset.

**Supplementary Table 5:** Significance of MsCr features across all data sets employing parametric statistics. *denotes p<0.01 **denotes p<0.001 ***denotes p<0.0001

| Data/measure | $a_\sigma$ | $b_\sigma$ | $a_\alpha$ | $b_\alpha$ | $a_\tau$ | $b_\tau$ | $a_\gamma$ | $b_\gamma$ | $a_\delta$ | $b_\delta$ | $a_{\sigma,\alpha}$ |
| --- | --- | --- | --- | --- | --- | --- | --- | --- | --- | --- | --- |
| DOC | *** |  |  |  |  | * |  | ** | ** |  |  |
| MCI |  | ** | ** | ** | *** |  | ** |  |  | * | *** |
| Schiz |  |  |  |  |  |  |  |  |  |  |  |
| Epilepsy | *** | ** |  |  |  | ** |  | *** | *** |  |  |
| Data/measure | $b_{\sigma,\alpha}$ | $a_{\sigma,\tau}$ | $b_{\sigma,\tau}$ | $a_{\sigma,\gamma}$ | $b_{\sigma,\gamma}$ | $a_{\sigma,\delta}$ | $b_{\sigma,\delta}$ | $a_{\alpha,\tau}$ | $b_{\alpha,\tau}$ | $a_{\alpha,\gamma}$ | $b_{\alpha,\gamma}$ |
| DOC |  |  | ** |  | * |  |  | * | * |  |  |
| MCI | *** | *** | ** | *** | ** | *** | ** | *** | *** |  |  |
| Schiz |  |  |  |  |  |  |  |  |  |  |  |
| Epilepsy |  | *** | *** | * | * | * |  |  |  |  |  |
| Data/measure | $a_{\alpha,\delta}$ | $b_{\alpha,\delta}$ | $a_{\tau,\gamma}$ | $b_{\tau,\gamma}$ | $a_{\tau,\delta}$ | $b_{\tau,\delta}$ | $a_{\gamma,\delta}$ | $b_{\gamma,\delta}$ | $\kappa$ | $\kappa_{gen}$ | $\alpha_{dropoff}$ |
| DOC | *** | *** |  |  | * | * |  |  |  | * |  |
| MCI |  |  |  |  | *** | *** |  | * |  | ** |  |
| Schiz |  |  |  |  |  |  |  |  |  |  |  |
| Epilepsy | *** | *** |  |  |  |  | *** | *** | *** |  | * |

**Supplementary Table 6:** Significance of MsCr features across all data sets employing non-parametric statistics. *denotes p<0.01 **denotes p<0.001 ***denotes p<0.0001

| Data/measure | $a_\sigma$ | $b_\sigma$ | $a_\alpha$ | $b_\alpha$ | $a_\tau$ | $b_\tau$ | $a_\gamma$ | $b_\gamma$ | $a_\delta$ | $b_\delta$ | $a_{\sigma,\alpha}$ |
| --- | --- | --- | --- | --- | --- | --- | --- | --- | --- | --- | --- |
| DOC | *** |  |  |  |  | * |  | ** | *** |  | * |
| MCI |  | * | * | * | ** |  |  |  |  |  | ** |
| Schiz |  | * |  |  | ** |  | * |  |  | *** |  |
| Epilepsy | *** | *** |  | ** |  | *** |  | *** | *** |  | *** |
| Data/measure | $b_{\sigma,\alpha}$ | $a_{\sigma,\tau}$ | $b_{\sigma,\tau}$ | $a_{\sigma,\gamma}$ | $b_{\sigma,\gamma}$ | $a_{\sigma,\delta}$ | $b_{\sigma,\delta}$ | $a_{\alpha,\tau}$ | $b_{\alpha,\tau}$ | $a_{\alpha,\gamma}$ | $b_{\alpha,\gamma}$ |
| DOC |  |  | * |  | * | * |  | * | * |  |  |
| MCI | ** | ** | * | ** | * | ** |  | * | * |  |  |
| Schiz |  |  |  |  |  |  |  | *** | *** | ** | ** |
| Epilepsy | *** | *** | *** | *** | *** | *** | *** |  |  |  |  |
| Data/measure | $a_{\alpha,\delta}$ | $b_{\alpha,\delta}$ | $a_{\tau,\gamma}$ | $b_{\tau,\gamma}$ | $a_{\tau,\delta}$ | $b_{\tau,\delta}$ | $a_{\gamma,\delta}$ | $b_{\gamma,\delta}$ | $\kappa$ | $\kappa_{gen}$ | $\alpha_{dropoff}$ |
| DOC | *** | *** |  |  | * | * | * | * |  | * |  |
| MCI |  |  |  |  | * | * |  |  |  | * |  |
| Schiz |  |  |  |  |  |  |  |  |  | * |  |
| Epilepsy | *** | *** | * |  | *** | *** | *** | *** | *** |  | *** |

